# LRRFIP2 modulates the response to hypoxia during embryonic cardiogenesis

**DOI:** 10.1101/2021.12.14.472540

**Authors:** Laura Ben Driss, Christophe Houbron, Florian Britto, Alain Schmitt, Morgane Le Gall, Philippe Daubas, Pascal Maire

**Affiliations:** Université de Paris, Institut Cochin, INSERM, CNRS. 75014 Paris, France; CNRS UMR 5048 - UM - INSERM U1054, 34090 Montpellier, France

## Abstract

Oxygen is crucial for appropriate embryonic and fetal development, including cardiogenesis. The heart is the first organ formed in the embryo and is required to provide oxygen and nutrients to all cells in the body. Embryonic cardiogenesis is a complex process finely regulated and prone to congenital malformations. It takes place in a hypoxic environment that activates the HIF-1α signaling pathway which mediates cellular and systemic adaptations to low oxygen levels. Since inhibition or overactivation of the HIF-1α signaling pathway in the myocardium lead to severe cardiac malformations and embryonic lethality, it is important that the cellular response to hypoxia be precisely regulated. While many gene regulatory networks involved in embryonic cardiogenesis have been characterized in detail, the modulation of the response of cardiomyocytes (CM) to hypoxia has remained less studied. We identified LRRFIP2 as a new negative cofactor of HIF-1α. Indeed, we have shown that the absence of *Lrrfip2* expression in a mouse KI model led to an enhance of many HIF-1α target genes including *Igfbp3, Bnip3* and *Ndufa4l2* in embryonic CM during development. As results, the absence of *Lrrfip2* led to the inhibition of the PI3K/Akt survival pathway, growth defects, mitochondrial dysfunction and to a precocious maturation of the embryonic CMs. Altogether, these defects led to the formation of a smaller heart unable to provide sufficient oxygen to the embryo and finally to a severe hypoxia and a precocious lethality.

**Graphical abstract:** 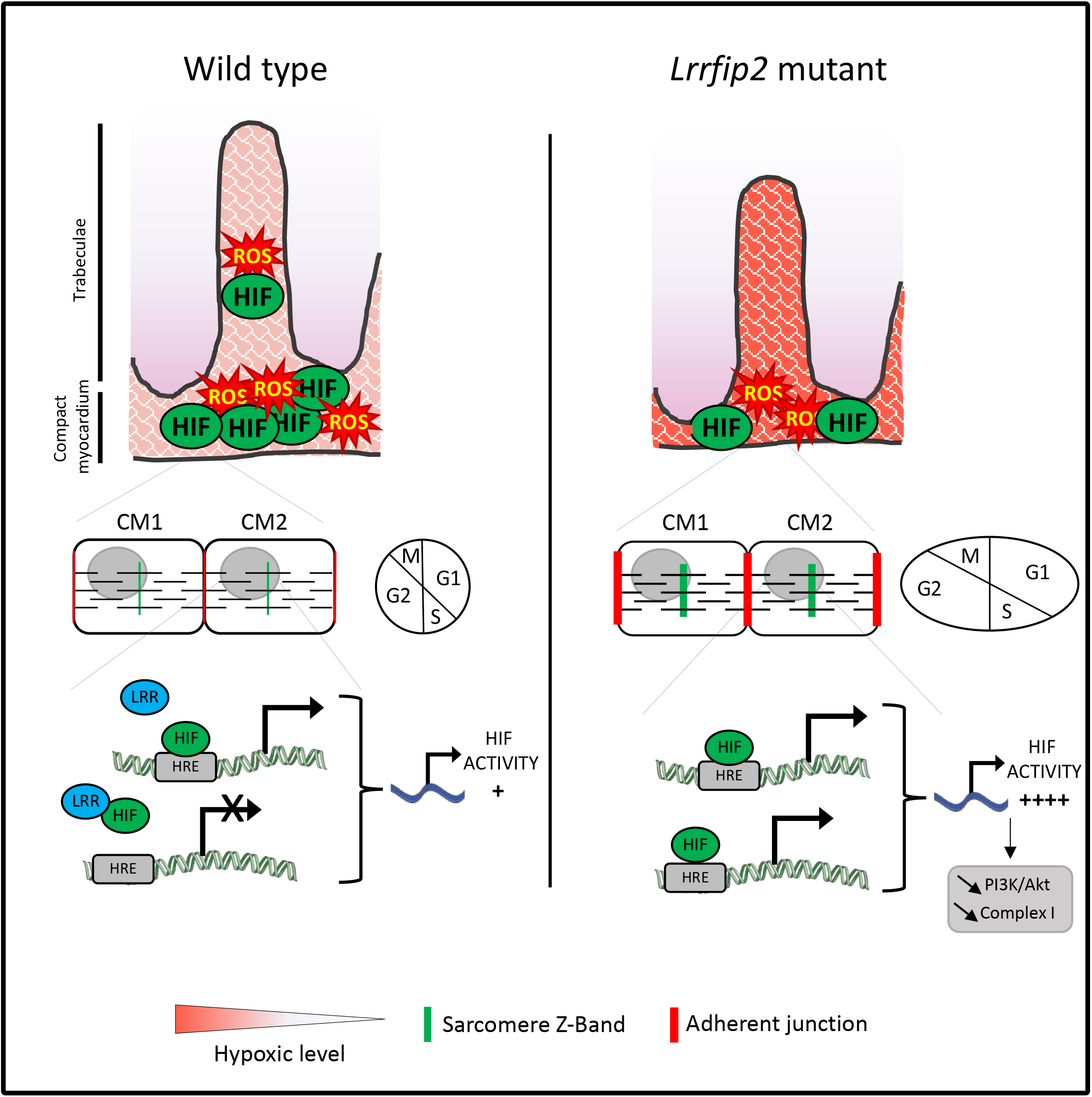

**Highlights:**

- LRRFIP2 regulates negatively the HIF-1α activity
- *Lrrfip2* deletion leads to an embryonic lethality between E11.5 and E13.5
- LRRFIP2 controls ROS production and CM maturation

## INTRODUCTION

The heart is the first organ to be formed during embryogenesis. It drives oxygen and nutrients to the other tissues and maintains vital functions during all life. Cardiogenesis is a complex process in which cardiac progenitors of different embryonic origins proliferate and differentiate into the diverse specialized cell types of the mature heart, including cardiomyocytes (CM) (Buckingham et al., 2005). Heart development is particularly prone to malformations and to congenital heart defects (CHDs) which affect 1% of live birth (Nicoll, 2018)(Houyel and Meilhac, 2021). These may be due to different combinations of genetic and environmental factors. One of the most important environmental risk during pregnancy is hypoxia (Ducsay et al., 2018). In case of low oxygen levels, Hypoxia Inducible Factors (HIFs) are induced. They control the expression of several genes implicated in the cellular and systemic adaptations to reduce oxygen availability through binding to their regulatory DNA regions. Under normoxia condition, HIF-1α is hydroxylated by the oxygen sensors prolyl hydroxylase (PHD), leading to its recognition by the von Hippel-Lindau (VHL) and to its degradation by the proteasome. Under hypoxia, HIF-1α escapes to this degradation and forms a heterodimer with HIF-1β/Arnt allowing the transcription of their target genes (Semenza, 2001).

In the embryonic heart, HIF-1α is the most expressed isoform of the HIFs gene family, its expression in CMs is associated with an anaerobic glycolytic metabolism (Menendez-Montes et al., 2016). During midgestation, the compact myocardium downregulates HIF-1α and CMs switch their metabolism toward an oxidative metabolism. Conditional HIF-1α knockout mice generated using *MLC2-Cre/Hif-1α-floxed* model leads to embryonic lethality at E11.5. This is associated with a strong increase of CMs proliferation leading to the absence of ventricular cavity formation and a break of cardiac morphogenesis (Krishnan et al., 2008). On another side, a specific deletion of *Vhl* in the myocardium using *Nkx2*.*5-Cre/Vhl-floxed* mice leads to HIF-1α stabilization and to embryonic lethality around E15.5-E17.5. This is the consequence of a decreased ventricular wall thickness and of a failure of the midgestational metabolic shift, impairing cardiac maturation and function (Menendez-Montes et al., 2016). So, the activation of HIF-1α in response to hypoxia is essential during cardiogenesis, but it is obvious that this activation has to be tightly regulated.

Embryonic cardiogenesis occurs in a hypoxic environment associated with a high ROS level in the CM until E9.0 (Krishnan et al., 2008). Then, ROS level progressively decreases leading to a decreased HIF-1α level required for the initiation of CM maturation. Therefore, embryonic hypoxia and ROS activate HIF-1α signaling pathway that is required for cardiac development. An uncontrolled increase or decrease in HIF-1α signaling during heart development drives to embryonic lethality.

Among the HIF-1α target genes, *Bnip3* (Feng et al., 2016) and *Igfbp3* (Huang et al., 2019) trigger mitochondria apoptosis and also autophagy and by this way facilitate programmed cell death. In fact, BNIP3 forms stable homodimer complexes inserted into mitochondrial membrane leading to damage and mitochondrial-dependent apoptosis. In another hand, IGFBP3 enhances apoptosis by sequestering IGF1 in the interstitial space preventing the activation of the IGF1R/PI3K/Akt survival pathway (Feng et al., 2016). *Ndufa4l2* is also activated by HIF-1α (Tello et al., 2011); it is an important element of the metabolic adaptation to hypoxia by reducing oxidative stress. In fact, NDUFA4L2 reduces intracellular ROS production under low oxygen conditions by negatively controlling the Complex I activity of the respiratory chain (Kruse et al., 2008). Overexpression of NUDUFA4L2 prevents also mPTP opening, cytochrome c release and Bax expression and by this way protects CMs against apoptosis (Kruse et al., 2008). So, in response to hypoxia, HIF-1α target genes are implicated in opposite functions promoting cell survival or in contrary cell death.

LRRFIP2 (Leucine Rich Repeat in Flightless Interacting Protein 2) is a protein expressed at a high level in the human heart. It is composed of two major domains by which it interacts with different partners among which Fightless (Fong and de Couet, 1999), Dishevelled (Liu et al., 2005), MYD88 (Dai et al., 2009) and NLRP3 (Jin et al., 2013). However, LRRFIP2 functions have been mostly studied in macrophages during the inflammatory process where it dampens the NLRP3 inflammasome activity (Jin et al., 2013)(Burger et al., 2016) but it was also characterized to activate the Wnt/ß-catenin pathway (Liu et al., 2005). Furthermore, *Lrrfip2* orthologue in Drosophila (CG8578) is required for correct sarcomere formation in indirect flight muscles (Schnorrer et al., 2010).

In this study, we show that LRRFIP2 interacts with HIF-1α and behaves as a negative modulator of the HIF-1α signaling pathway during heart development. We generated and analyzed a total knock-out of *Lrrfip2* and show that mutant embryos die around E11.5 due to numerous heart dysfunctions. LRRFIP2 mainly accumulated in the nuclear compartment of embryonic CM, appearing as a potential regulator of gene expression. Affymetrix transcriptomic analysis of E10.5 *Lrrfip2*^*-/-*^ and *Lrrfip2*^*+/+*^ hearts shows an overactivation of the HIF-1α signaling pathway confirmed by the overexpression of its target genes including *Igfbp3, Bnip3* and *Ndufa4l2*. Their overexpression leads to an inhibition of the PI3K/Akt/mTOR survival pathway, an inhibition of complex I of the respiratory chain associated with decreased ROS production and finally to a precocious maturation of mutant CMs.

## METHODS

### *Lrrfip2* GENE TARGETING VECTOR

We used the pDualvec targeting vector (Qu et al., 2006), allowing a unidirectional inactivation strategy. This targeting vector is composed by a positive selection marker, flanked by FLP recombinase recognition sequences and by the gene inactivation cassette. The gene inactivation cassette consists in a strong engrailed splice acceptor (SA) sequence, a reporter gene IRES-EGFP (Internal Ribosome Entry Site), and a polyadenylation signal sequence. This gene inactivation cassette is flanked on either side by oppositely oriented mutated LoxP sites. Once inserted into the intron of the target gene, it does not alter the expression of the target gene. Under the effect of a CRE recombinase, the inactivation cassette is definitively inverted, hijacking the transcriptional machinery via the splice acceptor, thus blocking expression of the target gene. Homologous DNA arms were amplified by PCR, using mouse 129/Sv genomic DNA as a template. To carry out the homologous arm subcloning, appropriate restriction sites were added in the forward and reverse primers. After subcloning of the 5’ homologous arm (XhoI-NotI, 3.1kb fragment), and the 3’ homologous arm (AscI-PmeI, 2.9kb fragment), amplified homologous fragments were sequenced to confirm absence of mutation that should have occurred during the amplification step. Linearized vector (25 μg) was electroporated into 129/Sv-derived embryonic stem (ES) cells (Kress et al., 1998). ES cells were selected with 150 μg/ml hygromycin 24 hours after electroporation. After 7 days of selection, 337 resistant ES cell clones were harvested and amplified separately **(Figure S1)**.

### ES CELL SCREENING AND GENERATION OF KNOCKOUT MICE

A rapid pre-screening was performed by PCR on both sides of the homologous regions, with an external primer outside of the targeting vector, and a primer into the gene inactivation cassette. Finally, 66 clones with a correct PCR profile were analyzed by Southern blot using external 5’ and 3’ probes with a SpeI digestion and an internal probe with a BamHI digestion.

Four of the clones contained a correct homologous recombination event. After karyotyping, two of the four ES cell clones were injected into C57BL/6 blastocysts. Chimeric males obtained were bred with C57BL/6 females to produce outbred F1 offspring carrying the modified *Lrrfip2-floxed-Hygro* allele. The *PGK-Hygro* selection cassette was removed to obtain *Lrrfip2*^*flox*^ allele by breeding *Lrrfip2-floxed-Hygro* mice with *FLPe* mouse strain (Rodriguez et al., 2000) to avoid potential influence on the expression of *Lrrfip2* gene. Afterwards, either *Lrrfip2*^*flox/+*^ mice were bred with *EIIa-Cre* animals expressing the Cre recombinase ubiquitously under the control of E2A promoter (Holzenberger et al., 2000) in order to obtain ubiquitous invalidation leading to *Lrrfip2*^*+/-*^ animals. Genotyping was carried out by PCR analysis on DNA extracted from tail samples. Detailed information for genotyping is available on request **(Figure S1)**.

### ANIMAL MODELS

Animal experimentations were carried out in strict accordance with the European STE 123 and the French national charter on the Ethics of Animal Experimentation. Protocols were approved by the Ethical Committee of Animal Experiments of the Institut Cochin, CNRS UMR 8104, INSERM U1016. The *Lrrfip2*^*+/-*^ mouse mutant line was maintained on a C57Bl6/N genetic background. As *Lrrfip2*^*-/-*^ animals die during embryogenesis, mutant embryos are generated by crossing *Lrrfip2*^*+/-*^. Fetuses were staged, taking the appearance of the vaginal plug as E0.5, and harvested at the indicated day. The yolk sac or tail of embryos was used for DNA extraction and genotyping by PCR using allele-specific primers.

### LRRFIP2-WT-FW

5’TTCCATGAGTTAGGGTTCTTCC; **LRRFIP2-WT-RV**: 5’GTTCTGAGAGCAACACTGGTC; **LRRFIP2-KO-FW**: 5’TTCCATGAGTTAGGGTTCTTCC; **LRRFIP2-KO-RV**: 5’CATCGTCATCCTTGTAGTCCTCG. Hearts from E9.5 or E10.5 embryos were isolated and either frozen in liquid nitrogen for RNA or proteins extractions, or used for primary cell culture. Embryos were isolated and fixed in 0.4% paraformaldehyde two hours at room temperature, rinsed in PBS and kept in 20% sucrose-PBS for two hours and embedded in 10-10% gelatin/sucrose-PBS, frozen in cold isopentane (−30°C) and sectioned on a cryostat (10µm). Hypoxyprobe (HPI #HP1-100) was injected (60mg/kg) in pregnant mice two hours before sacrifice. Embryos were collected and included as previously described, the slides were post-fixed with acetone (+4°C) for 10minutes and then practiced as described in the “immunofluorescence” section.

### CELL CULTURE

Primary cultures were derived from hearts of E10.5 mice embryos. *Lrrfip2*^*+/+*^ and *Lrrfip2*^*-/-*^ hearts were mechanically dissociated and cultured on DMEM-Glutamax (Gibco #31966-021), 5% FBS, 10% SVF on well previously coated with gelatin at 37°C in an incubator with 5% CO2 and 95% air. H9c2 cells were grown in DMEM (Gibco #41965-039) including 10% (v/v) fetal bovine serum, 1% (v/v) penicillin-streptomycin solution (v/v) at 37°C in an incubator with 5% CO2 and 95% air. Under hypoxia, primary CMs and H9c2 cells were cultured in an incubator with 10% CO2 and 1% O2 for varying times.

### RECOMBINANT VIRUS TRANSDUCTION IN PRIMARY CARDIOMYOCYTE CELL CULTURE

To induce in culture the inversion of the floxed *Lrrfip2* allele in CM culture, *Lrrfip2*^*flox/flox*^ CM were transduced twice with Ad-mGFP (Vector Biolabs #1060) or Ad-Cre-mGFP (Vector Biolabs #1779) adenoviruses (200 MOI) and then kept in culture during three days before analysis.

### CELL TREATMENTS

To activate the HIF-1α signaling pathway, H9c2 cells were cultured with 200µM of CoCl2 (Sigma #C8661) during 30min, 4h or 24h. To stimulate ROS generation, H9c2 cells were cultured with 20µM and 100µM of Rotenone (Sigma #R8875) during 30min, 4h or 24h. Rotenone was diluted into dimethyl sulfoxide (DMSO).

### DIHYDROETHIDIUM (DHE) STAINING

Primary CM and H9c2 cells were cultured on 24 wells plates. After quick removal of the culture medium, cells were washed once with PBS, then incubated for 8 minutes at RT with a solution of 3µM DHE (Thermo Fisher #D11347). Nuclei are stained with Hoechst. Cells are further washed with PBS and immediately observed. Fluorescent image acquisition was performed using the Zeiss Axiovert 200M microscope.

### siRNA AND cDNA TRANSFECTION

To generate a LOF model, primary CM and H9c2 cells were transfected with short interfering RNA (siRNA) at 10nM using Lipofectamine RNAimax (Invitrogen) 24h and 48h after plating and analyzed at 96h. Control (Sigma #SIC001), and a mix of two *Lrrfip2* (Sigma #s89557 #89558) siRNAs were used. The efficiency of the interference was controlled by RT-qPCR and WB. To generate a GOF model, primary CM and H9c2 cells were transfected with plasmids at 100nM using Lipofectamine 300 (Invitrogen), 24h after plating and analyzed at 72h. LRRFIP2-Δ1 and LRRFIP2-FL in pCR3™ expression vectors were used and the efficiency of transfection was estimated by RT-qPCR and WB.

### RT-qPCR

RNA was extracted in TRIzol (Thermo Fisher Scientific) from cell cultures or isolated hearts. RNAs were reverse-transcribed using the Superscript III kit (Invitrogen) according to the manufacturer’s instructions. Reverse transcription was performed with 100 ng of RNA. Quantitative PCR analysis was performed using a Light Cycler 480 (Roche) according to the manufacturer’s instructions using a SYBR Green I kit (Roche #04887352001). Values were normalized to the expression level of housekeeping genes *36B4* or *18S*. The primers used are referenced on Table1.

### AFFYMETRIX MICROARRAYS

Affymetrix analysis were performed from embryonic hearts at E10.5 distributed into two groups: WT (n=3) and KO (n=2). Total RNAs were obtained using TRIzol reagent and DNAse1 treatment (Quiagen). RNA integrity was certified on bioanalyzer (Agilent). Hybridization to Mouse Gene 2.0-ST arrays (Affymetrix) and scans (GCS3000 7G expression Console software) were performed on the Genom’IC plateform (Institut Cochin, Paris). Probe data normalization and gene expression levels were processed using the Robust Multi-array Average (RMA) algorithm in expression Console software (Affymetrix). Gene ontology analysis was performed using Ingenuity (IPA) and GSEA softwares.

### IMMUNOFLUORESCENCE

Embryo heart sections are blocked 1h at RT in PBS-Triton (15% horse serum, 0.5% Triton X-100). Immunofluorescence was performed with a standard protocol, using primary antibodies, listed in Table2, overnight at 4°C, Alexa Fluor conjugated secondary antibodies (Invitrogen 1:1000) and Hoechst nuclear staining, during 1h at RT. Heart sections were finally mounted in Mowiol mounting medium and kept at 4°C until image acquisition.

### IMAGE ACQUISITION

Fluorescence images were acquired using Olympus BX63F microscope, Zeiss Axiovert 200M microscope or Spinning-Disk IXplore. Images were merged and edited with ImageJ. Background was reduced using brightness and contrast adjustments applied to the whole image.

### IMMUNOPRECIPITATION AND WESTERN BLOT ANALYSIS

Embryonic hearts and cells were lysed in RIPA buffer (Sigma #R0278). Immunoprecipitation of protein extracts was performed using a standard protocol based on magnetic beads coupled to bacterial protein G (Invitrogen #10003D). 1µg of anti-LRRFIP2 antibodies per 500µg of protein extracts was used. Proteins were eluted in Laemmli 1x buffer.

Embryonic hearts and cells were lysed in RIPA buffer (Sigma #R0278) and proteins were separated through denaturating SDS-PAGE electrophoresis using Mini-Protean TGX precast gels 4-15% (Biorad) and transferred on Nitrocellulose (0.2μm, Biorad) membrane using the Trans-Blot turbo transfer system (Biorad). Membranes were blocked with 5% milk-TBS-1% Tween (TBST) 1 hour at room temperature and probed overnight at 4°C with primary antibodies in 5% milk-TBST-1X, listed in Table3. Membranes were then hybridized with secondary antibodies coupled to HRP (ThermoFisher 1:10 000). Proteins were revealed using SuperSignal West Femto substrate (ThermoFisher).

### ELECTRON MICROSCOPY

E10.5 embryonic hearts were dissected in cold ADS-1X/0,5%Glucose and fixed with 2,5% glutaraldehyde + 4% PFA-PBS. After one hour, the samples were put in a fresh solution until analysis.

### STATISTICS

Quantitative data sets were analyzed using an unpaired non-parametric Mann Whitney test, two-way ANOVA with Tukey’s multiple comparisons test, using GraphPad Prism6.0 software. Statistical significance was set at a pvalue<0.05.

## RESULTS

### *Lrrfip2* is ubiquitously expressed in adult mice

Except its functions in macrophages during the inflammation process, few is known concerning the expression pattern and functions of LRRFIP2 in adult mice. By RT-qPCR, we showed that *Lrrfip2* mRNA was detected in all tested tissues as already observed in human. It accumulated mostly in the heart and skeletal muscles and at a lower level in the stomach (Figure 1a). Western blot confirmed the accumulation of LRRFIP2 proteins in different adult tissues and revealed several isoforms of which the largest 110kDa is detected specifically in the heart while other isoforms around 55kDa accumulated in all tissues including the heart (Figure 1b).

**Fig. 1.**
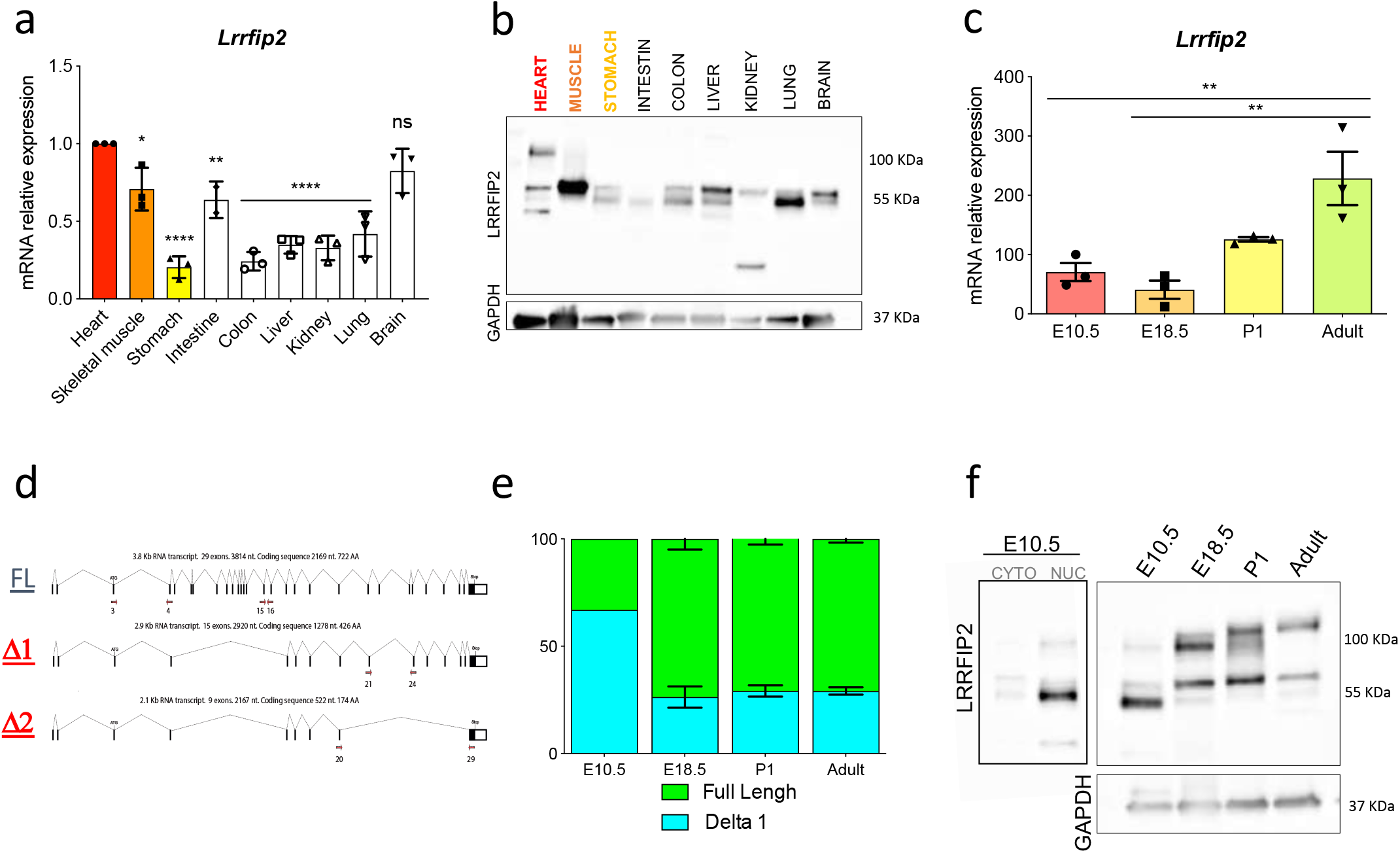
*Lrrfip2* is strongly expressed in the heart and its protein accumulates in the nucleus of embryonic CM. **a**. Quantification of *Lrrfip2* gene expression in different indicated adult tissues. **b**. LRRFIP2 isoforms in adult tissues by Western blot experiments with GAPDH control expression. **c**. Quantification of *Lrrfip2* in the heart from embryonic to adult cardiogenesis. **d**. Identification of 3 alternative *Lrrfip2* mRNAs in E10.5 embryonic heart by PCR experiments. **e**. PCR and **f**. Western Blot experiments showing the preferential accumulation of LRRFIP2 in the nuclear fraction (NUC) as compared with the cytoplasmic fraction (CYTO) and a switch in the relative quantity of the different LRRFIP2 isoforms during heart development. Two way anova significance test (prism software) : *p-value<0.05 **p-value<0.01 ****p-value<0.001.

### *Lrrfip2* expression is temporally regulated in developing heart

We further characterized the expression pattern of *Lrrfip2* in the embryonic and adult heart by RT-qPCR and observed an increased mRNA accumulation from Embryonic day 10.5 (E10.5) to the adult stage (Figure 1c). With specific primers located along *Lrrfip2* cDNA, we identified three different transcripts expressed in E10.5 heart. The first transcript, called full length (FL), is composed by all *Lrrfip2* exons leading to the 110kDa identified protein and two alternative transcripts: Δ1 with a splicing event between the exons 4-17 coding for a 55kDa protein and Δ2 spliced between the exons 4-17 and 20-29 coding for a 25kDa protein (Figure 1d). By PCR experiments, we further determined the relative proportion of these different transcripts during embryonic and adult cardiogenesis. We focused at four different crucial stages of heart development: early cardiogenesis (E10.5), just before birth (E18.5), during CMs binucleation process (post-natal day 1, P1) and in the adult when CMs are post-mitotic cells unable to proliferate. We observed at E10.5, a ratio of 75% Δ1 for 25% of FL. From E18.5 this ratio was inverted in favor of the FL transcripts (75/25%). Whatever the developmental stage, the Δ2 isoform was very minor compared to the two other isoforms (Figure 1e). Accordingly, at E10.5 we observed a small protein around 55kDa accumulating preferentially in CMs nuclei (Figure 1f). From E18.5, a higher isoform around 110kDa accumulated with the 55kDa protein, according with mRNAs isoforms expression, and finally, at the adult stage, the FL protein became the more abundant isoform in the heart (Figure1f). LRRFIP2 is thus temporally regulated during cardiogenesis and presented different isoforms according to the stage of life.

Altogether our results showed that *Lrrfip2* expression is controlled by transcriptional and post-transcriptional mechanisms during cardiogenesis.

### *Lrrfip2* deletion causes structural cardiac defects and embryonic lethality

To determine the role of LRRFIP2 during heart development, we performed its invalidation in ES cells by introducing a LoxP cassette composed of a strong splicing acceptor site (SA) associated with an Ires-EGFP reporter and a Flp-PGK-Hygromycin selection cassette. After ES cells selection, two separate KI heterozygous mouse lines were crossed with an EIIa-Flipase to specifically delete the PGK-Hygromycin cassette. After its excision, the two mouse *Lrrfip2*^*flox/+*^ lines were further crossed with an EIIa-CRE line (Holzenberger et al., 2000) to give rise to *Lrrfip2*^*+/-*^ animals that were viable and fertile. After the recombination, the LoxP-cassette is inverted leading to a premature STOP codon and to the absence of *Lrrfip2* transcription (Figure 2a). We confirmed the absence of *Lrrfip2* mRNAs in *Lrrfip2*^*-/-*^ embryos at E10.5 by RT-qPCR with specific primers of the WT and recombined allele (REC) and by Western blot with specific antibodies able to recognize all LRRFIP2 isoforms (Figure 2b). With the two independent lines, we never obtained newborn *Lrrfip2*^*-/-*^ animals suggesting an embryonic lethality. In fact, we determined that all *Lrrfip2*^*-/-*^ embryos were dead after E13.5 and we observed a loss of the Mendelian ratio for live embryos after E11.5 suggesting the onset of their lethality at this stage (Figure 2c). By MF20 immunostaining, we showed that *Lrrfip2*^*-/-*^ hearts were smaller with thin ventricular walls and defective trabeculae from E10.5 to E13.5, while no obvious phenotype was observed at E9.5 (Figure 2d, e). A significant reduced CMs number was detected in *Lrrfip2*^*-/-*^ hearts, which is neither associated with defects in CM number per µm2 nor in CM area (Figure 2f-h). The smaller heart observed in E10.5 mutant embryos was therefore the consequence of its reduced number of CMs as compared with E10.5 WT hearts. Surprisingly we observed no difference in the proportion of Ki67+/MF20+ CM or in cyclin and check-point markers expression in the cell cycle in WT and mutant embryos (Figure 2i, j and Figure S2) but a significant decreased number of pHH3+/MF20+ CM in the mutant (Figure 2k, l). These results suggested an important defect in the proliferative capacity of mutant CMs and showed that LRRFIP2 is essential for embryonic CM proliferation and heart chambers development.

**Fig. 2.**
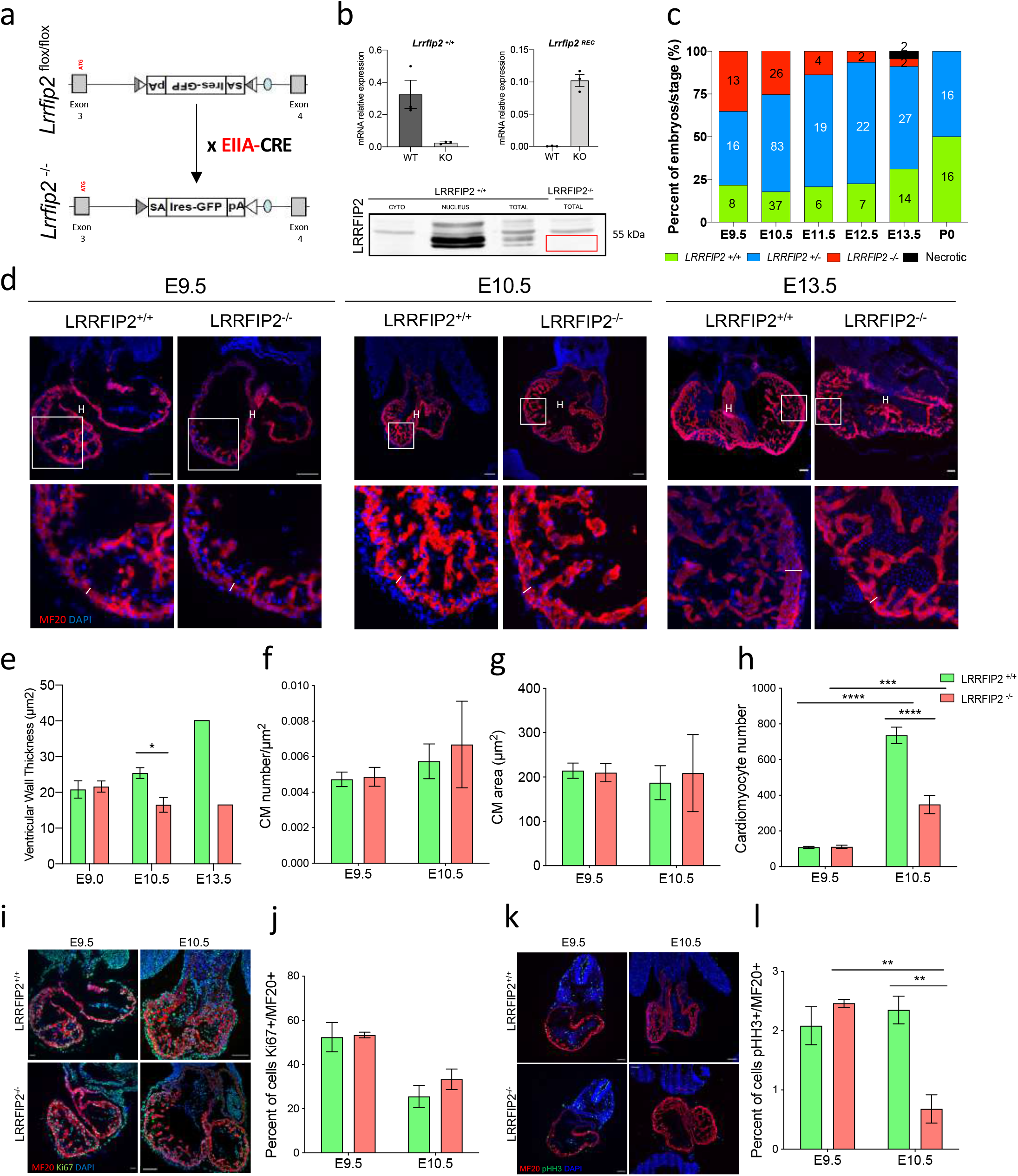
*Lrrfip2* is required for myocardial development. **a**. *Lrrfip2* Flox allele before and after EIIA-CRE recombination. **b**. Validation by RT-qPCR and WB of *Lrrfip2* WT and inverted allele in E10.5 embryos. **c**. Histogram of the distribution and number of *Lrrfip2*^*+/+*^, *Lrrfip2*^*+/-*^ and *Lrrfip2*^*-/-*^ embryos at different stages indicating the death of *Lrrfip2*^*-/-*^ embryos between E11.5 and E13.5. **d.e** Quantitative immunofluorescence on *Lrrfip2*^*+/+*^ and *Lrrfip2*^*-/-*^ heart sections of E9.5, E10.5 and E13.5 embryos showing smaller heart and thin ventricular walls in mutant embryos. **f**. Quantification of CMs number per µm2 and **g**. CMs area of E9.5 and E10.5 *Lrrfip2*^*+/+*^ and *Lrrfip2*^*-/-*^ hearts. **h**. Histogram showing the number of CM present in 9.5 and E10.5 *Lrrfip2*^*+/+*^ and *Lrrfip2*^*-/-*^ hearts. **i**. Ki67 immunofluorescence on heart sections and **j**. % of Ki67^+^/MF20^+^ CMs in E9.5 and E10.5 *Lrrfip2*^*+/+*^ and *Lrrfip2*^*-/-*^ embryos. **k**. pHH3 immunofluorescence on heart sections and **l**. % of pHH3^+^/MF20^+^ CM in E9.5 and E10.5 *Lrrfip2*^*+/+*^ and *Lrrfip2*^*-/-*^embryos. Two-way anova significance test (prism software): *p-value<0.05 **p-value<0.01 ****p-value<0.001 – Scale bar 100µm. H: heart. The white lines indicate the thickness of ventricular walls.

### LRRFIP2 modulates HIF-1α signaling pathway

To better understand how the absence of *Lrrfip2* leads to midgestational lethality, we performed Affymetrix transcriptomic analysis of WT and mutant E10.5 hearts. We detected 80 genes downregulated and 142 genes upregulated more than 1.5-fold in mutant hearts as compared with WT hearts (p<0.05). With IPA software, we identified HIF-1α, IL-1ß and ATF4 as upstream transcriptional regulators predicted to be activated (Figure 3a). We focused more on the HIF-1α pathway in the rest of our study considering its role during cardiogenesis. According to Affymetrix data, we observed that the expression of *Hif-1*α was not controlled by *Lrrfip2* (Figure 3b) but interestingly we showed a significant decrease of the accumulation of HIF-1α in the heart in the absence of LRRFIP2 (Figure 3c,d). Analysis of Affymetrix data showed an increase of HIF-1α target genes expression that we confirmed by RT-qPCR on WT and mutant E10.5 hearts (Figure 3e,f). Altogether, these data showed that absence of *Lrrfip2* during cardiogenesis leads to an increased expression of many HIF-1α target genes suggesting that the role of LRRFIP2 in embryonic CMs may be to dampen HIF-1α activity.

**Fig. 3.**
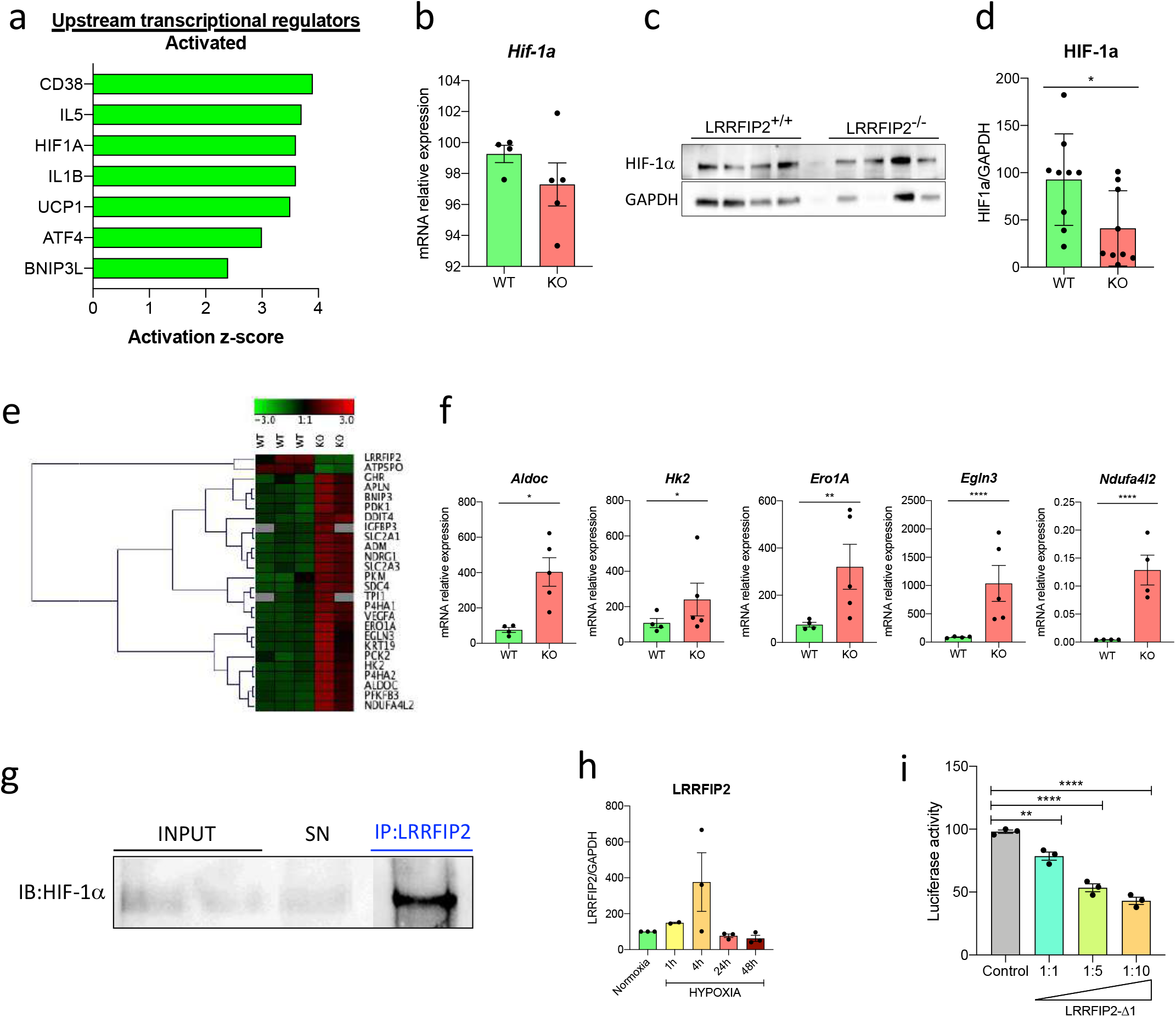
LRRFIP2 modulates HIF-1α signaling pathway. **a**. IPA analysis and upstream activated transcriptional regulators (positive z-score) from Affymetrix data of mRNAs extracted from *Lrrfip2*^+/+^ and *Lrrfip2*^*-*/-^ E10.5 hearts. **b-d**. RT-qPCR and Western Blot analysis of *Lrrfip2*^*+/+*^ (n=4) and *Lrrfip2*^*-/-*^ (n=4) 10.5 hearts showing that the absence of LRRFIP2 has no effect on *Hif-1*α mRNA accumulation but leads to a decreased HIF-1α protein accumulation in mutant hearts. **e**. Heat Map from Affymetrix data representing up and down-regulated genes of the HIF-1α signaling pathway in *Lrrfip2*^+/+^ and *Lrrfip2*^*-*/-^ E10.5 hearts. **f**. HIF-1α target genes are upregulated in E10.5 *Lrrfip2*^*-/-*^ as compared with WT hearts as estimated by RT-qPCR. Data were normalized by 36B4 expression. **g**. Western blot revealed with HIF-1α antibodies on E10.5 total protein extracts of WT hearts immunoprecipitated (IP) with LRRFIP2 antibodies, input and IP supernatant (SN) were used as controls. **h-i**. H9c2 cells were challenged with Hypoxia, Cocl2 or Rotenone treatments for the indicated time. *Lrrfip2* mRNA and LRRFIP2 protein levels were estimated by RT-qPCR and WB and normalized by *18S* and GAPDH respectively. t-test significance test (prism software): *p-value<0.05 **p-value<0.01 ****p-value<0.001

To better understand how LRRFIP2 could modulate the HIF-1α signaling pathway, we first tested a potential direct interaction between these two proteins in WT E10.5 heart by immunoprecipitation (IP) assays. After an IP with LRRFIP2 antibodies, we were able to reveal HIF-1α by western blot, demonstrating an interaction between these two proteins during *in vivo* cardiogenesis (Figure 3g). We also challenged H9c2 cells with different treatments known to modulate HIF-1α stability and determined LRRFIP2 expression after hypoxia, CoCl2 and Rotenone treatments during very short (1h), short (4h) and long term (24h) processing. However, 4h of hypoxia led to an increased LRRFIP2 accumulation (Figure 3h) while CoCl2 and Rotenone treatments had no really effect on LRRFIP2 stability (Figure S3). To assess if LRRFIP2 can directly modulate the HIF-1α/HRE activity, we performed luciferase assay with a 6x-HRE reporter transfected in H9c2 cells under hypoxia. We used increasing concentration of LRRFIP2-Δ1 and we observed a dose-dependent decrease of HRE activity in response to LRRFIP2 (Figure 3i). Altogether, these results identified LRRFIP2 as a new negative cofactor of HIF-1α.

### LRRFIP2 modulates the hypoxia signaling pathway during embryogenesis

Since we observed an overactivation of HIF-1α signaling in E10.5 mutant hearts (Figure 3e, f), we looked for the origin of this activation. As HIF-1α is activated in response to hypoxia, we first determined the proportion of hypoxic tissues in embryos with pimonidazole, an hypoxyprobe. Pimonidazole incorporates in hypoxic cells below a P_O2_ of 10 mm Hg and is further labeled with MAb1 antibodies. To assess the oxygenation status of mutant embryos, we injected the pimonidazole in pregnant mice two hours before their sacrifice (Figure 4a). At E9.0, when the embryos were growing under hypoxia as physiological conditions, we observed that most of WT and mutant embryonic cells were hypoxic, with increased MAb1 staining in the mutant embryos. At E10.5 on the contrary we observed a strong increase of hypoxic tissues in mutant embryos and more specifically in their heart as compared with WT embryos where MAb1 staining decreased between E9 and E10.5 (Figure 4b,c). As hypoxia is the result of a decreased O_2_ availability, we investigated the state of the vascular system. We observed an increased expression of the HIF-1α target gene *Vegfa* in mutant hearts but we did not detect vascular defects in E10.5 mutant embryos by CD31 labeling (Figure 4d,e). Altogether these results suggested a role of LRRFIP2 in the regulation of hypoxia during embryogenesis independent of the vascularization process.

**Fig. 4.**
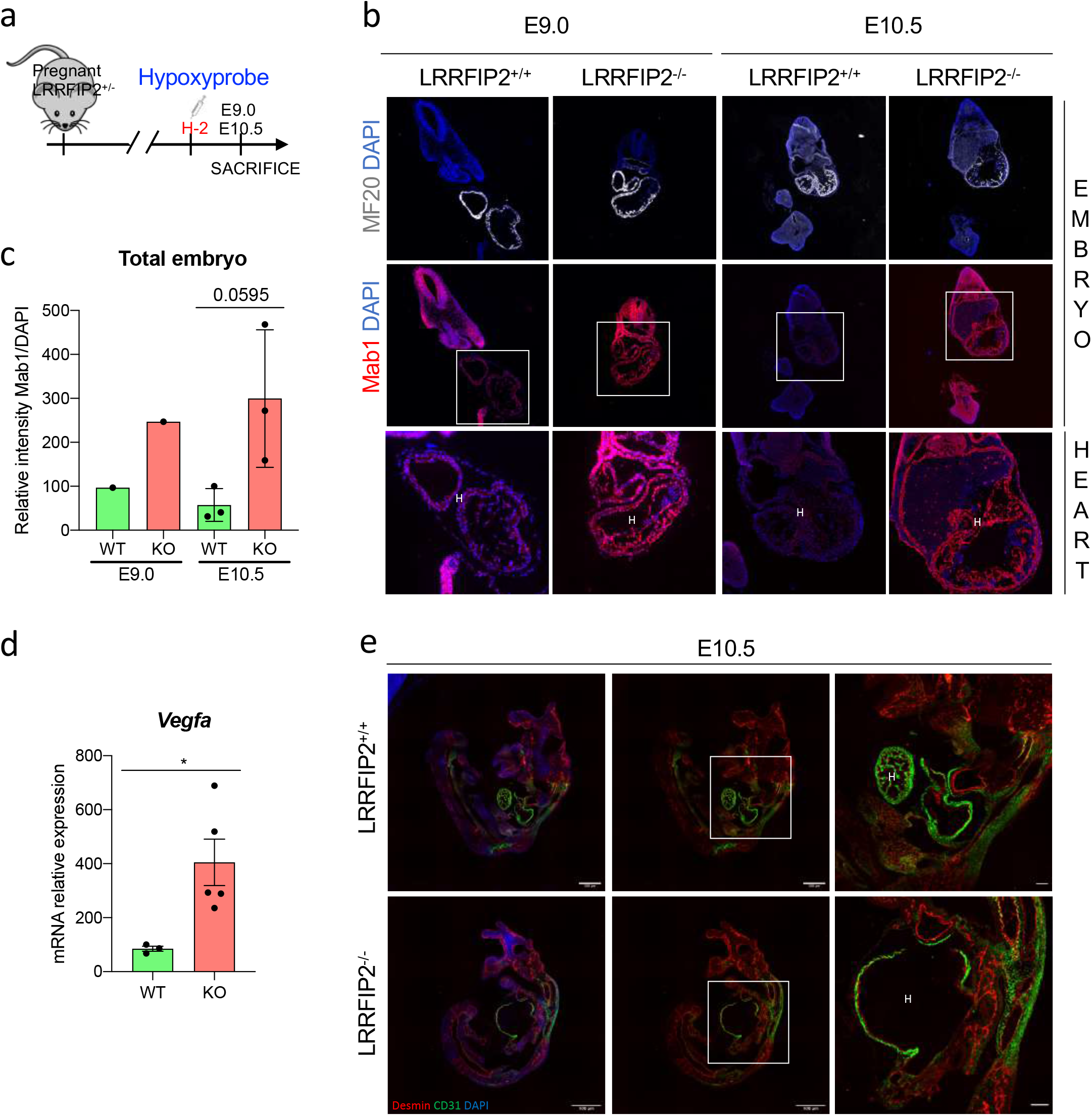
LRRFIP2 controls hypoxia during embryogenesis. **a**. The hypoxyprobe pimonidazole was injected in E9.0 and E10.5 pregnant females 2 hours before their sacrifice and revealed by MAb1 antibody. **b.c**. CMs were revealed by MF20 staining and MAb1^+^ cells revealed on *Lrrfip2*^*+/+*^ and *Lrrfip2*^*-/-*^ embryo sections at the heart level. Zoom of the heart is presented. Quantification of MAb1 staining in E9.0 and E10.5 *Lrrfip2*^*+/+*^ and *Lrrfip2*^*-/-*^ hearts. Absence of LRRFIP2 expression leads to a strong increase of hypoxia from E9.0 to E10.5 in the entire embryo. **d**. Increased *Vegfα* expression in E10.5 *Lrrfip2*^*-/-*^ hearts estimated by RT-qPCR. **e**. CD31 and Desmin staining at the heart level in *Lrrfip2*^*+/+*^ and *Lrrfip2*^*-/-*^ E10.5 embryos. t-test and Two way anova significance test (prism software) : *p-value<0.05 **p-value<0.01 ****p-value<0.001 – Scale bar 100µm. H: heart.

### Ectopic activation of the HIF-1α signaling pathway disrupts CM survival pathways and favors their death

HIF-1α regulates the expression of genes implicated in several processes like cell survival and cell death. Among those genes, *Igfbp3* inhibits the PI3K/Akt survival pathway and *Bnip3* is involved in apoptosis and autophagy. We confirmed their overexpression in mutant hearts by RT-qPCR experiments (Figure 5a,d). As a consequence of *Igfbp3* and *Redd1* overexpression, we observed a strong decrease of the phosphorylation level of Akt(S473) and of its targets S6K and 4-eBP1 in E10.5 mutant hearts showing a severe inhibition of this survival pathway in the absence of LRRFIP2 (Figure 5b,c). We also observed a tendency to an increased Caspase3+ CMs number in mutant hearts suggesting an increased apoptosis of mutant CMs (Figure 5e,f). Altogether, our results show that during cardiac embryogenesis *Lrrfip2* dampens HIF-1α signaling leading to the downregulation of the PI3K/Akt cell survival pathway and to a decreased protection against apoptosis.

**Fig. 5.**
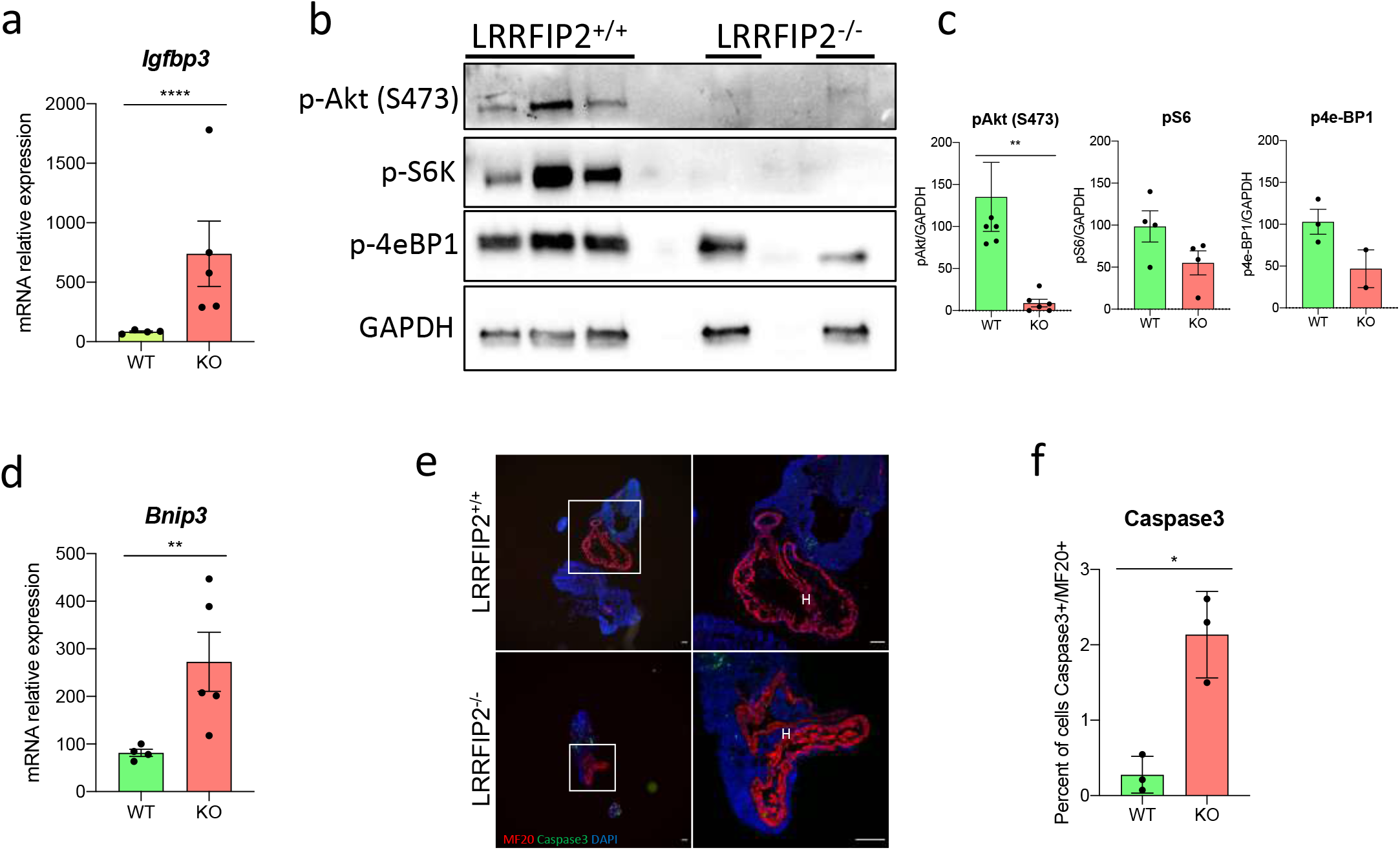
Ectopic activation of HIF-1α signaling pathway disrupts CMs survival pathways and favors their death. **a**. RT-qPCR on *Lrrfip2*^*+/+*^ and *Lrrfip2*^*-/-*^ 10.5 hearts showing the upregulation of *Igfbp3* expression in mutant samples. **b.c**. WB quantification of proteins from *Lrrfip2*^*+/+*^ and *Lrrfip2*^*-/-*^ E10.5 hearts showing decreased phosphorylation of Akt(S473), S6K and 4eBPB1. **d**. RT-qPCR on WT and mutant E10.5 hearts showing the overexpression of *Bnip3* in mutant samples. **e.f**. Immunostaining quantification of Caspase3^+^/MF20^+^ CMs on *Lrrfip2*^*+/+*^ and *Lrrfip2*^*-/-*^ 10.5 heart sections, and corresponding percentage of Caspase3^+^/MF20^+^ CMs. t-test significance test (prism software) : *p-value<0.05 **p-value<0.01 ****p-value<0.001 – Scale bar 100µm. H: heart.

### HIF-1α overactivation upon *Lrrfip2* deletion leads to mitochondrial dysfunction

HIF-1α also regulates the expression of genes involved in cell metabolism and mitochondrial functions. In the Affymetrix transcriptomic data previously described, we identified “mitochondrial dysfunction” and “oxidative phosphorylation” in the top three of the canonical pathways impacted by the absence of LRRFIP2 (Figure 6a). To validate these data, we first quantified the amount of mitochondrial (mt)DNA in comparison of nuclear (n)DNA in control and mutant E10.5 hearts. By qPCR experiments we observed no difference in mtDNA content in 10.5 mutant heart, suggesting that the same quantity of mitochondria accumulated in WT and mutant CMs (Figure 6c). By electron microscopy (EM) in contrast, we identified two populations of mitochondria in E10.5 mutant hearts: a population with small mitochondria with an area comparable to that observed in WT hearts and another population made up of larger mitochondria with a swelling aspect (Figure 6d,e). On the whole, genes involved in the oxidative phosphorylation process are downregulated in *Lrrfip2*^*-/-*^ E10.5 hearts (Figure 6b) excepted *Ndufa4l2* known to inhibit the Complex I of the respiratory chain, which was upregulated (Figure 3f). We observed a strong accumulation of NDUFA4L2 in all the myocardium in mutant hearts compared to their control (Figure 6g) suggesting increased inhibition of the respiratory chain in mutant CMs and consequently a decreased ROS production. We thus measured ROS accumulation in WT and mutant E10.5 hearts to test this possibility. Accordingly, by DHE staining on primary cultures of WT and mutant CM, we showed a decreased ROS level in mutant cells (Figure 6h). These results suggested that LRRFIP2 controlled mitochondrial activity during heart development.

**Fig. 6.**
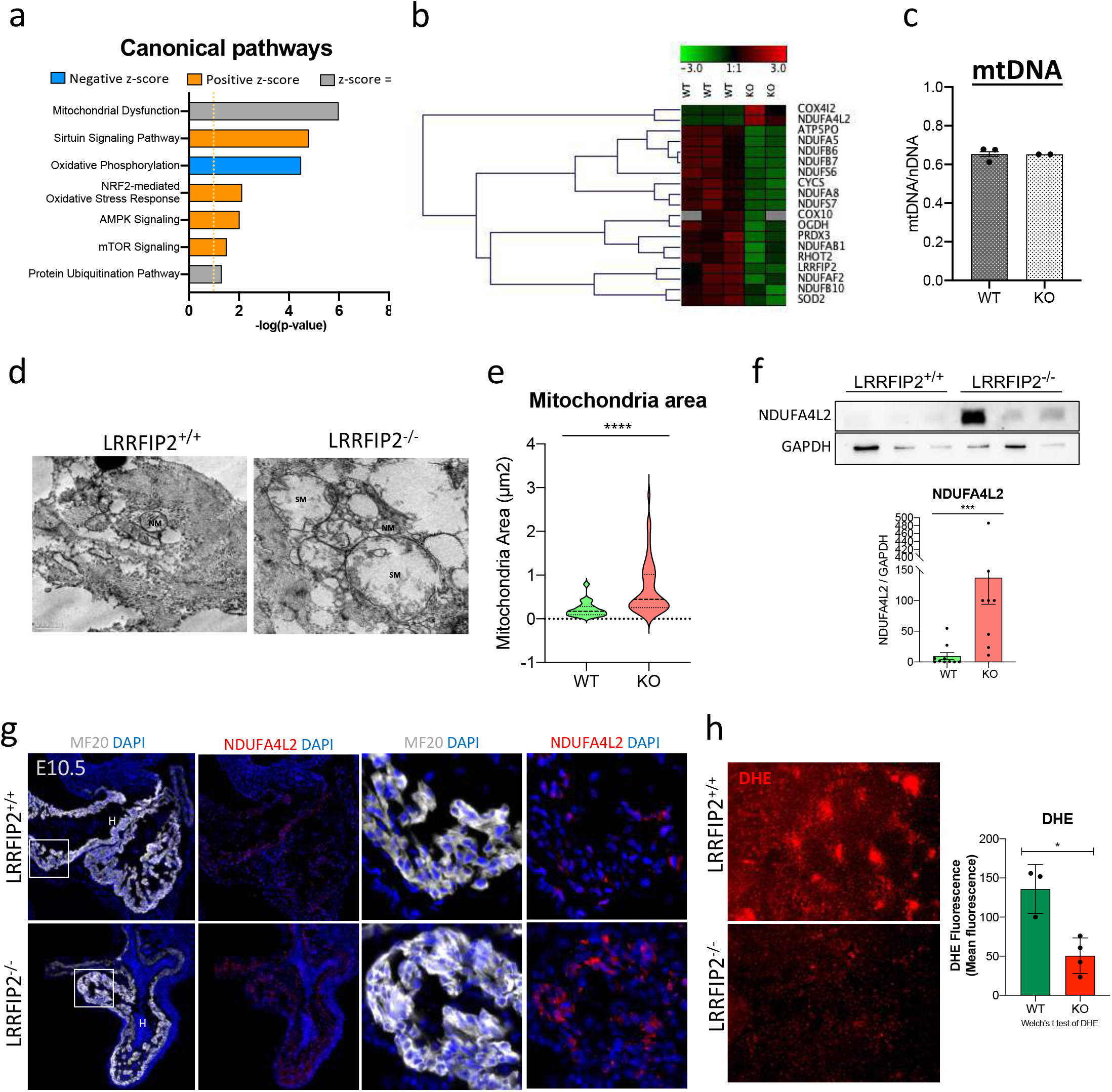
HIF-1α overactivation leads to mitochondrial dysfunctions. **a**. IPA analysis of Affymetrix *Lrrfip2*^*+/+*^ and *Lrrfip2*^*-/-*^ 10.5 hearts identified deregulated canonical pathways among which “oxidative phosphorylation” appearing as inhibited. **b**. Heat Map representation from Affymetrix data showing a global inhibition of genes with mitochondrial functions excepted *Cox4i2* and *Ndufa4l2*. **c**. Quantification of mtDNA/nDNA ratio in DNA from *Lrrfip2*^*+/+*^ and *Lrrfip2*^*-/-*^ 10.5 hearts **d.e**. Electron microscopy on *Lrrfip2*^*+/+*^ and *Lrrfip2*^*-/-*^ 10.5 heart sections identified two mitochondria populations in *Lrrfip2*^*-/-*^ hearts, one with increased area only present in mutant samples. **f**. WB quantification and **g**. immunostaining showing a strong accumulation of NDUFA4L2 in *Lrrfip2*^*-/-*^ (n=3) as compared with *Lrrfip2*^*+/+*^ (n=3) E10.5 hearts. **h**. DHE quantification showing decreased ROS levels in E10.5 *Lrrfip2*^*-/-*^ CM primary cultures. t-test significance test (prism software) : *p-value<0.05 **p-value<0.01 ****p-value<0.001 – Scale bar 100µm. NM: normal mitochondria. SM: swelling mitochondria.

### Premature decreased ROS level leads to a precocious maturation of CMs

Reduction of ROS level is a signal for CMs maturation initiation during embryonic cardiogenesis (Murray et al., 2013). As we observed a decreased ROS levels in mutant CMs (Figure 6h), we investigated the maturation state of WT and mutant CMs. In E10.5 *Lrrfip2*^*-/-*^ hearts, we observed a precocious formation of sarcomeres by immunostaining (α-Actinin and phalloidin) and by EM, as compared with WT samples (Figure 7a,b). Sarcomeres of mutant CMs appeared more differentiated and better organized. They also presented a slight increase of the Z-Band length and a higher Z-band thickness (Figure 7c,d). Cell-cell junctions also participate to the CM maturation. By western blot, we did not detect any defect in N-Cadherin quantity in mutant as compared to WT hearts, but by immunostaining we showed that N-Cadherin was more compact between mutant CMs, and we observed bigger junctions by EM (Figure 7e,f). So, differences in the organization of the adherent junctions were obvious between mutant and WT CMs. By EM, we also observed bigger desmosomes, involved in mechanical force transmissions, between mutant CMs (Figure 7g). Altogether these results suggested that LRRFIP2 slowed down the maturation and favored the proliferation of CMs during embryonic cardiogenesis through several complementary mechanisms.

**Fig. 7.**
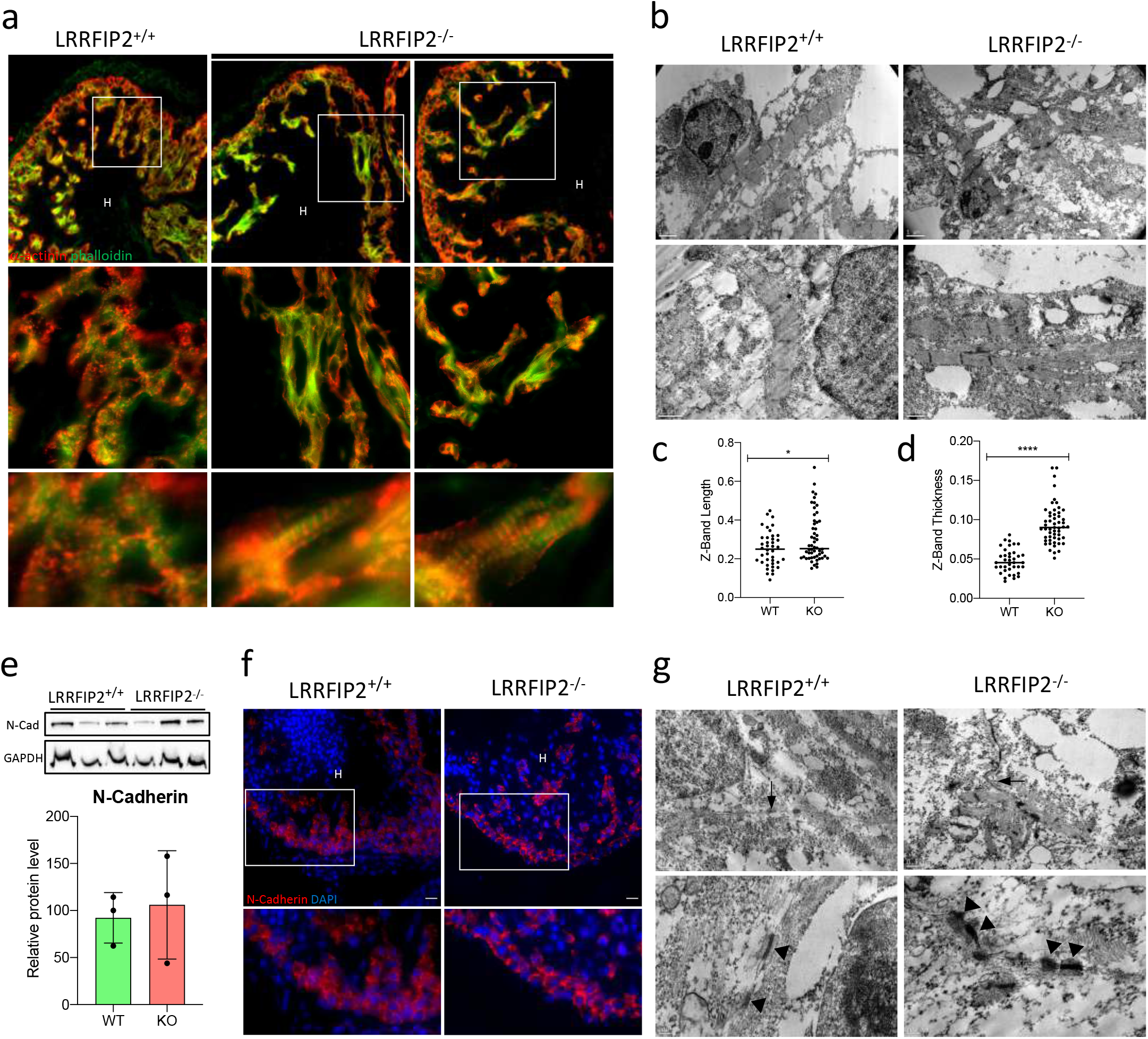
*Lrrfip2* deletion leads to a premature maturation of CM. **a**. Immunostaining of α-Actinin and phalloidin and **b**. electronic microscopy showing a precocious maturation of sarcomeres of *Lrrfip2*^*-/-*^ 10.5 hearts. **c.d**. Z-bands length and thickness in sarcomeres of *Lrrfip2*^*+/+*^ and *Lrrfip2*^*-/-*^ 10.5 hearts. **e**. WB quantification of N-Cadherin in *Lrrfip2*^*+/+*^ and *Lrrfip2*^*-/-*^ E10.5 hearts **f**. Immunostaining highlighted N-Cadherin compaction at the junctions between two mutant cells. **g**. Electronic microscopy showing bigger junctions (arrows) and bigger desmosomes (head of arrows) between CM of mutant E10.5 hearts. t-test significance test (prism software) : *p-value<0.05 **p-value<0.01 ****p-value<0.001 – Scale bar 100µm

**Fig. 8.**
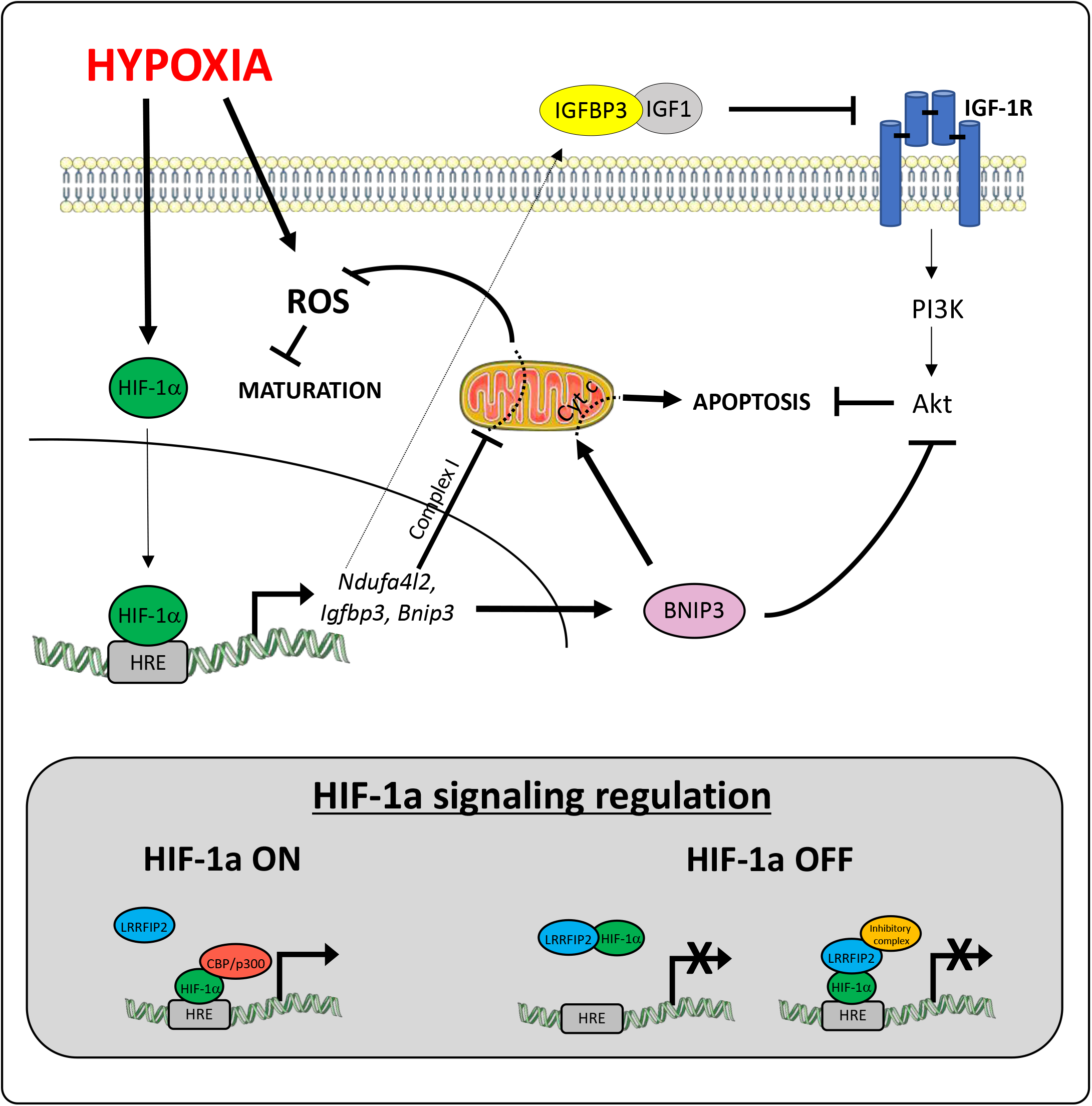
Fig. 8. Proposed model of HIF-1α signaling regulation by LRRFIP2. In the absence of LRRFIP2, HIF-1α activity is increased leading to an overexpression of different target genes like *Ndufa4l2, Bnip3* or *Igfbp3*. These transcriptional enhance cause an inhibition of the survival pathway PI3K/Akt, promoting apoptosis and ROS decrease. All together, this phenotype leads to a precocious maturation, a smaller heart and a strong hypoxia. The HIF-1α signaling can be regulated by a direct interaction between HIF-1α and LRRFIP2.

## DISCUSSION

Heart development occurs in a hypoxic environment resulting in the activation of the HIF-1α signaling pathway. Both Cardiac specific loss of function and overactivation of this pathway during embryogenesis lead to cardiac hypoplasia and hyperplasia respectively, which are responsible of their embryonic precocious lethality (Krishnan et al., 2008; Menendez-Montes et al., 2016). We focused our study on LRRFIP2, a protein ubiquitously expressed with a higher expression in the heart (Fong and de Couet, 1999). *Lrrfip2* deletion in the embryo led to embryonic lethality at mid-gestation associated with significant cardiac malformations such as cardiac hypoplasia observed from E10.5. We characterized LRRFIP2 as a new partner of HIF-1α and we showed that this interaction dampened the HIF-1α activity. *Lrrfip2* mutant CM presented increased HIF-1α transcriptional activity leading to a reduced proliferation and increased maturation of the CM. Finally, LRRFIP2 appeared as a new negative HIF-1α cofactor.

### LRRFIP2 MODULATES HIF-1α SIGNALING PATHWAY AND HYPOXIA

Loss of function of Lrrfip2 in the mouse embryo led to embryonic lethality from E11.5 onwards associated with significant cardiac malformations such as decreased ventricular wall thickness and a smaller heart due to a reduced number of CMs. This phenotype is very close to the one observed in *Vhl*-mutant in which the decreased CM number observed is the consequence of the overactivation of the HIF-1α signaling pathway (Menendez-Montes et al., 2016). Transcriptomic analysis performed on embryonic hearts at E10.5 also identified an overactivation of the HIF-1α signaling pathway in the absence of LRRFIP2. We identified an interaction between LRRFIP2/HIF-1α in the heart and a nuclear localization of LRRFIP2 in CM suggesting that it may directly modulate the HIF-1α transcriptional activity. Accordingly, the expression of different HIF-1α target genes, described as deregulated in the transcriptomic analysis like *Bnip3, Igfbp3, Vegfa, Ero1a, Aldoc* or *Ndufa4l2*, showed that the absence of LRRFIP2 led to an overexpression of the HIF-1α target genes. Based on these observations, LRRFIP2 appears as a negative regulator of the HIF-1α signalization during heart development.

During cardiogenesis, hypoxia is associated with the activation of the HIF-1α signaling and is maintained up to E9.0 from when a progressive reduction of its activation is detected. In the absence of LRRFIP2, we observed a maintained level of hypoxia in mutant embryos that did not appear to be the consequence of vascularization defects, as estimated by CD31 labeling, but by altered cardiac properties.

### LRRFIP2 PROMOTES CMs SURVIVAL AND PROLIFERATION

Overactivation of the HIF-1α signaling pathway caused by the absence of LRRFIP2, led to the overexpression of *Igfbp3* known to sequester IGF1 in the intermembrane space. This sequestration impairs downstream PI3K/Akt activation that is involved in cell growth and survival pathway (Huang et al., 2019). Accordingly, we observed an important inhibition of the PI3K/Akt pathway in E10.5 *Lrrfip2*^*-/-*^ embryos as estimated by Akt and S6 protein phosphorylations. Moreover, *Bnip3*, another HIF-1α target gene known to be pro-apoptotic, is also overexpressed (Huang et al., 2019). The increased number of Caspase3^+^ CM observed in *Lrrfip2*^*-/-*^ embryos at E10.5 may be the consequence of this increase *Bnip3* expression or to mitochondrial defects. Finally, inhibition of the PI3K/Akt survival pathway and pro-apoptotic signaling activated in the absence of LRRFIP2 present LRRFIP2 as a protector against CM cell death during cardiogenesis.

Severe mitochondrial defects were also suggested by the transcriptomic analysis, and were associated with a decreased expression of genes implicated in the oxidative phosphorylation pathway (OXPHOS). No major modification was noticed in the mtDNA quantity present in E10.5 mutant CM, but an important defect in their size was observed. In fact, mitochondria appeared bigger with a swelling aspect in mutant CM suggesting important mitochondrial dysfunction (Kwong and Molkentin, 2015). This pathological aspect observed at E10.5 in *Lrrfip2*^*-/-*^ CM probably participates to the absence of heart growth observed. It will be important to determine the number of Caspase3^+^ mutant CM at E11.5 to test this hypothesis. While many genes implicated in the oxidative phosphorylation pathway are down-regulated in mutant CM, *Ndufa4l2* known to inhibit the complex I of the respiratory chain (Li et al., 2017) is overexpressed. NDUFA4L2 accumulated in all myocardium of *Lrrfip2*^*-/-*^ embryos at E10.5 and may participate to the lower ROS level observed in mutant embryos as a consequence of decreased complex I activity.

Finally, we also observed an early maturation of E10.5 mutant CMs in terms of sarcomerization, junction thickness and proliferative capacity. This early maturation may be due to decreased in ROS production in mutant CMs, because a decrease in ROS level is a signal for maturation initiation (Murray et al., 2013). Our results identify LRRFIP2 as a factor promoting survival and CM proliferation rather than maturation by acting on modulation of HIF-1α target gene expression.

## CONCLUSION

In this study, we identified LRRFIP2 as a new negative cofactor of HIF-1α acting during embryogenesis to control heart development. While the precise mechanisms of LRRFIP2 to dampen HIF1-α activity remains to be clarified, we showed that heart growth is blunted from E10.5 in *Lrrfip2*^*-/-*^ embryos associated with an upregulation of numerous genes by HIF-1α. *Lrrfip2*^*-/-*^ embryos died between E11.5-E13.5, well before *Vhl*^*-/-*^ embryos. That could be explained by the fact that in the case of *Lrrfip2*, the deletion occurs in the entire embryo while *Vhl* is only deleted in the myocardium in the mutant embryo (Menendez-Montes et al., 2016) or that other signaling pathways than HIF-1α, may be affected in *Lrrfip2*^*-/-*^ embryos. LRRFIP2 is known to modulate Wnt/ß-Catenin, NLRP3, NFΚB and Flightless/Actin signaling pathways. The activity of these pathways may be impaired in *Lrrfip2*^*-/-*^ embryo and participate in the identified phenotype, although our transcriptomic analysis did not favor this hypothesis. Our work showed that LRRFIP2 is mainly involved during mouse embryonic development to control heart growth through interaction with HIF-1α. As *Lrrfip2* is expressed in many adult tissues including heart, it may also participate in the control of several pathophysiological processes involving hypoxia such as regeneration after myocardial ischemia (Zhang et al., 2019).

## Supporting information

Sup Figures Ben Driss

## ACKNOWLEDGMENTS

We thank Athanassia Sotiropoulos, Fanny Bajolle, Sigolène Meilhac and Carole Peyssonnaux for helpful discussions.

We thank Cochin IMAG’IC and GENOM’IC platforms for helpful advice.

L.B.D was supported successively by a PhD fellowship from the Fondation pour la recherche médicale (FRM, DPC20171239475) and by the Agence Nationale pour la Recherche (ANR, MYOLINC -16-CE14-0032-01). Financial support to this work was provided by the FRM (DPC20171239475), the Association Française against Myopathy (AFM n°20766), the Institut National de la Santé et de la Recherche Médicale (INSERM) and the Centre National de la Recherche Scientifique (CNRS).

Designed experiments, L.B.D, C.H, F.B, P.D, P.M. Performed experiments, L.B.D, F.B, C.H, A.S. Interpreted data, L.B.D, F.B, C.H, P.D, P.M. Bioinformatic analysis, M.L.G. Wrote the manuscript, L.B.D, P.M with the input of F.B. Funding Acquisition, P.M, P.D.

