## Supplementary material for "LRRFIP2 modulates the response to hypoxia during embryonic cardiogenesis": Sup Figures Ben Driss

### Figure S1

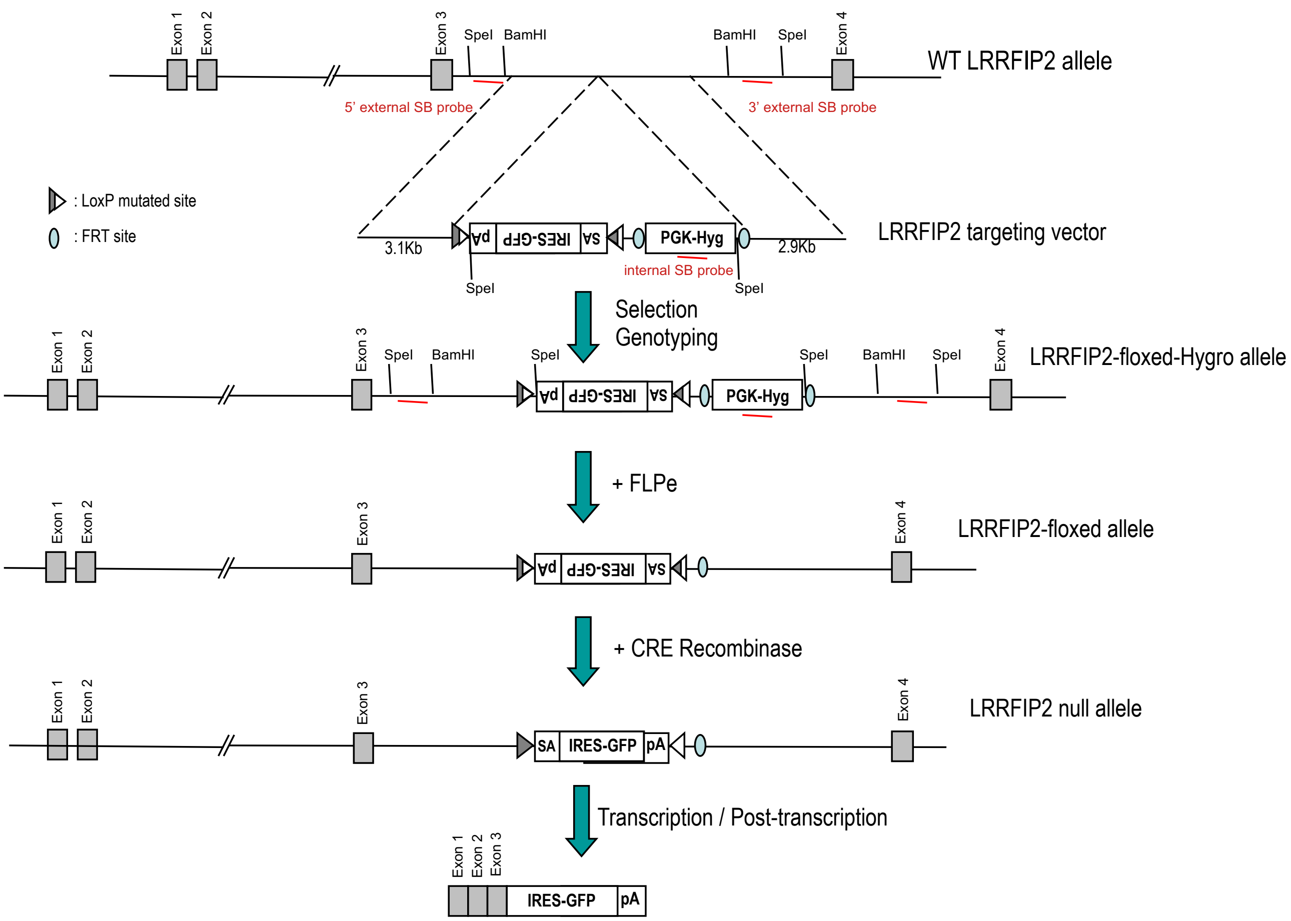


**Figure Supp 1 :** *Lrrfip2* Flox allele before and after EIIA-CRE recombination

### Figure S2

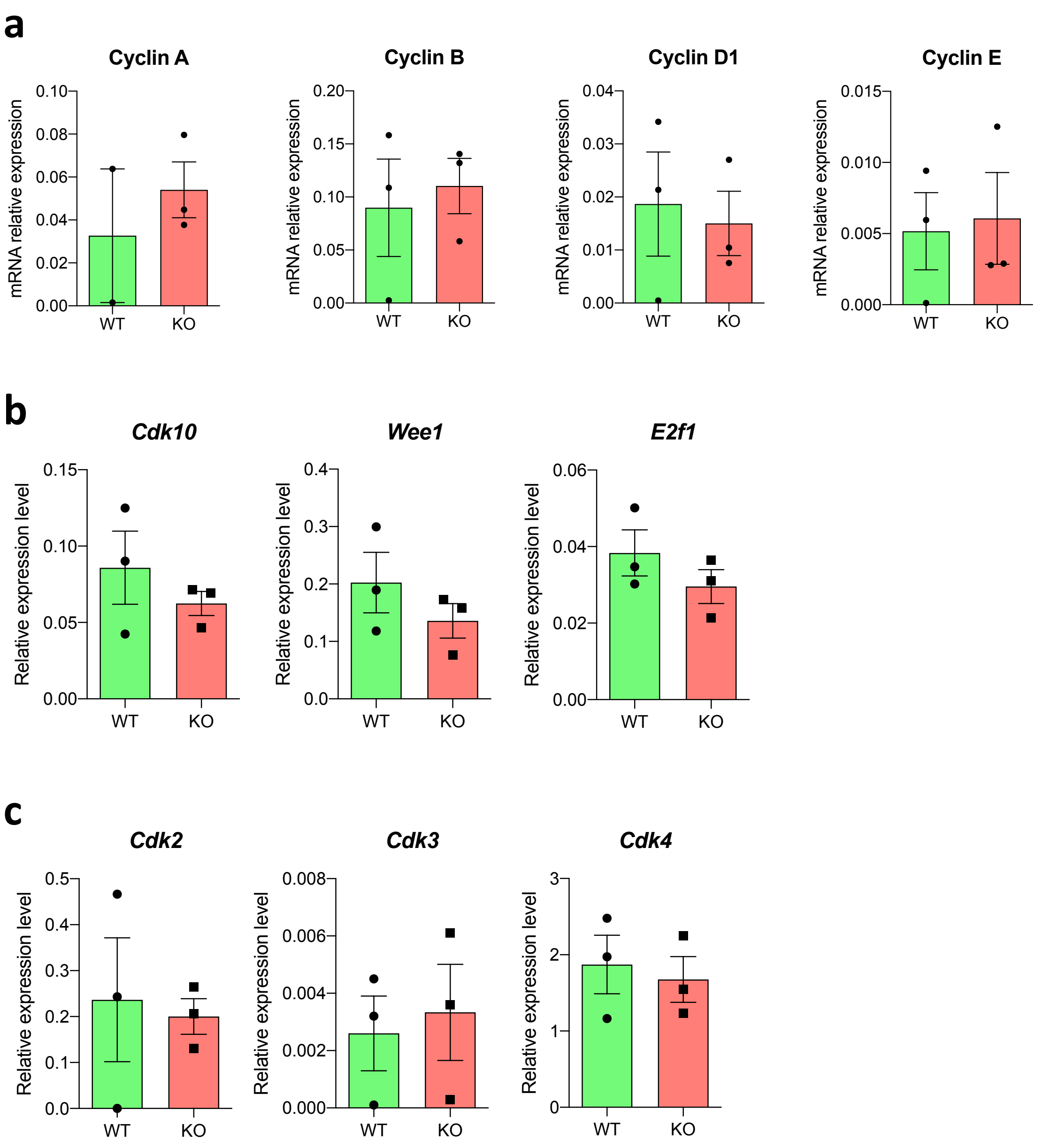


**<u>Figure Supp 2 :</u>** mRNA expression of different cell cycle markers by Rt-qPCR from E10.5 hearts. **a.** mRNA expression of cyclins. **b.** mRNA expression of genes implicated in the G1/S checkpoint. **c.** mRNA expression of genes implicated in the G2/M checkpoint. Normalized by 36B4.

**Figure S3**

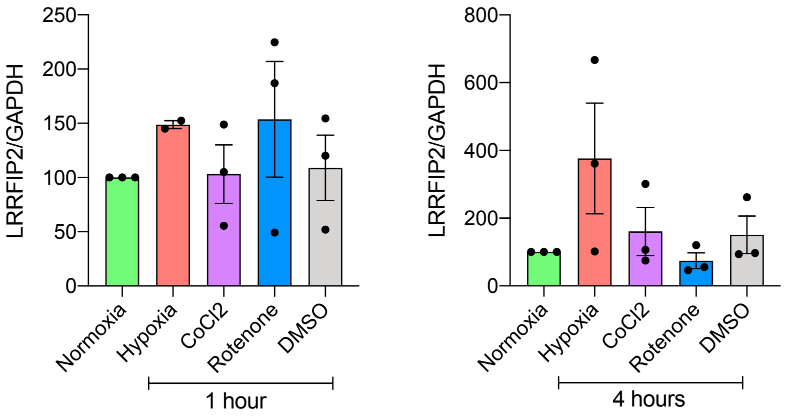


**<u>Figure Supp 3 :</u>** H9c2 cells were challenged with Hypoxia, Cocl2 or Rotenone treatments for the indicated time. LRRFIP2 protein levels were estimated by WB and normalized by GAPDH.
